# A versatile high throughput strategy for cloning the env gene of HIV-1

**DOI:** 10.1101/2020.04.01.020735

**Authors:** Nitesh Mishra, Ayushman Dobhal, Shaifali Sharma, Kalpana Luthra

## Abstract

The trimeric envelope glycoprotein (gp120/gp41)_3_ of human immunodeficiency virus-1 (HIV-1) mediates viral and host cell membrane fusion, initiated by binding of viral envelope gp120 protein to the CD4 receptor on host immune cells. Functional env genes from infected individuals have been widely used as templates for vaccine design, for setting up viral neutralization assays and to study the viral evolution and pathogenesis. Traditional topoisomerase or T4 DNA polymerase mediated approaches for cloning single genome amplified (SGA) env genes are labor-intensive, cost-ineffective with low-throughput, thereby enabling functional analysis of only a limited number of env genes from the diverse circulating quasispecies in infected individuals. Herein, we report an efficient, easy to optimize and high-throughput approach for cloning diverse HIV-1 env genes. Multiple env/rev gene cassettes, derived from infected infants, were subjected to SGA using Phusion polymerase and utilized as megaprimers in overlap extension PCR mediated cloning (OEC), circumventing the requirement for novel enzymes. Furthermore, utilization of Phusion polymerase for both the amplification of env/rev cassettes and OEC allows convenient monitoring and optimization, thereby providing much greater flexibility and versatility for analysis of env genes from HIV-1 infected individuals.

## Introduction

Rapid evolution of HIV-1 strains is increasing the global viral diversity. High baseline rates of mutation (nucleotide substitution rate) due to an error prone reverse transcriptase, rapid replication cycle, in combination with continuous immune-driven selection within the host generates a unique and complex viral quasispecies in each infected individual (1,2). In the context of HIV-1, the viral species in each infected individual accrues immunologically relevant mutations, generating a viral pool of highly complex and unique variants circulating within each host (3–5). Population based consensus viral sequences are used to define wildtype HIV-1 virus, but given the extensive diversity, each variant within a clade is distant from the consensus sequence, giving rise to millions of coexisting variants in circulation. Developing an HIV-1 vaccine is a global priority, and the discovery of broadly neutralizing antibodies (bnAbs) has invigorated the field of HIV-1 vaccine research (6,7). bnAbs target the envelope glycoprotein (env) of HIV-1 within defined epitopes, though inferred germlines (iGLs) of several bnAbs do not bind to most of the difficult-to-neutralize HIV-1 isolate. Identification and/or engineering of env variants capable of engaging and shepherding bnAb development pathway are a key focus of lineage-based vaccine approach (8–12).

Functional characterization of env gene provides key information for vaccine design, identify env candidates capable of serving as immunogens, understanding viral evolution and pathogenesis in response to host response, or mutational landscape among several other virological and immunological functions (3–5,8,13,14). Traditionally, envelopes are characterized by generating pseudoviruses (env genes from different hosts cloned in mammalian expression vectors) in combination with env deficient HIV-1 backbones (such as PSG3∆env, ZM247∆env) (15–18). Conventional approaches for amplifying env gene via single genome amplification (SGA) and cloning env gene are time and labor intensive processes. In addition, utilization of topoisomerases (TOPO cloning) or T4 DNA polymerases (SLIC, or Infusion cloning) to clone env gene can only be done at low throughout as increasing cost associated with cloning multiple env genes restricts the cloning to few select env clones (15,19,20). Single env clones are not representative of the diversity of HIV-1 in a patient’s blood sample (19,21–25). The SGA technique using traditional TOPO or Infusion cloning kits to generate functional env clones that represent majority of the circulating quasispecies is low-throughput and not cost-effective due to the involvement of multiple PCR reactions, purification steps and ligation reactions.

In the age of high-throughput molecular and systems biology, several approaches have been described to overcome the limitations associated with conventional cloning approaches (26–41). Genes are often identified as difficult-to-clone that exhibit features such as uneven base distribution, secondary structures, toxicity to bacterial strain used for cloning and the HIV-1 env gene exhibits all of these features (42–44). Few select strategies exist by which the env gene can be cloned with high efficiency. One such approach called overlap extension cloning (OEC) involves solely a a PCR-based cloning workflow that offers highest versatility in terms of optimizing cloning approaches (26–28). In OEC, the insert of interest (called a megaprimer) is used in conjunction with a proofreading polymerase that does not have strand displacement activity (such as Phusion polymerase) to swap the insert into vector backbone, and is sequence and ligation independent. Variants of OEC like IOEP (Improved Overlap Extension PCR) (29), IVA (In Vivo Assembly) (39), AQUA (Advanced QUick Assembly) (40), CPEC (Circular Polymerase Extension Cloning) (41) have initiated new avenues for high-throughput cloning. Most of these approaches work well for subcloning (swapping inserts from one plasmid backbone into another) but fail when applied to cloning gene of interest from biological samples. This study aimed to design a cost-effective and high-throughput approach to clone HIV-1 env gene in order to capture maximum functional diversity of the circulating viral variants in infected individuals. A low-cost, easy to optimize, and high-throughput method, based on the principles of overlap extension cloning, to construct env clones representing the functional diversity of circulating HIV-1 viral variants is proposed herein, allowing rapid production of heterogenous patient derived env genes. The robustness of this strategy was further evaluated and confirmed by cloning viral env gene from eight HIV-1 infected infants.

## Results and Discussion

Overlap extension cloning (**Fig. 1**) is a PCR-based cloning technique and therefore primer design is critical for a successful OEC, as non-specific primer binding can lead to amplification of spurious PCR products. Conserved regions of the HIV-1 envelope were selected for primers design to clone the viral env gene, to accommodate for the high envelope sequence diversity, uneven base composition (on average 36% adenine, 22% thymine, 24% guanine and 18% cytosine) and high propensity of secondary structure in the AT-rich HIV-1 genome (42,43). Given that HIV-1 diverges overtime (1,2), we first updated our primer repository to ensure optimum amplification of the env gene (onwards called env/rev cassettes, as partial fragment for rev gene is part of the env gene). Two-step nested PCR reactions were used to amplify env/rev cassettes, where the forward primers (Fw1 and Fw2) bind to the tat region (upstream to the env gene), and the reverse primers (Rv1 and Rv2) bind to nef region (downstream to the env gene) (**Fig. 2a**). Our major goal was to design a cloning strategy where env/rev cassettes can be cloned in a single tube without the need for novel enzymes like topoisomerases, exonucleases, T4 DNA polymerases, Taq DNA ligase or recombinases. Therefore, the robustness of the nested PCR for env/rev amplification was of utmost importance. Using the parameters of Clade C tat and nef sequences of Indian origin available in Los Alamos National Laboratory’s HIV Database, we selected a total of 276 tat (HXB2 numbering 5831 – 6045) and 140 nef (HXB2 numbering 8797 – 9417) sequences. Sequences were aligned with HIValign tool using the HMM-align model with compensating mutation occurring within 5 codons to compensate frameshift and using previously reported primers (15–17) as anchors [VIF1 as Fw1 primer (HXB2 numbering 5852 – 5876), ENVA as Fw2 primer (HXB2 numbering 5951 – 5980), OFM19 as Rv1 primer (HXB2 numbering 9604 – 9632) and ENVN as Rv2 primer (HXB2 numbering 9144 – 9172)] (**Fig. 2a**). A total of 4 Fw1, 2 Rv1, 12 Fw2 and 6 Rv2 primers were selected (**Table 1**) to address the diversity observed within the geographically restricted sequences of Indian origin (**Fig. 2b – c**). We first utilized a multiplex PCR approach where all Fw1 and Rv1 primers were pooled for the first round, and all Fw2 and Rv2 primers were pooled for the amplification via nested PCR. Though consistent amplification of env/rev cassettes across samples was achieved, the efficiency of PCR was low and in addition smearing and/or non-specific amplification were notably observed (**Fig. 2d**). To overcome this, we next performed nested PCR using all of the possible primer pair combinations (a total of 576 individual PCR reactions), Four of these primer pairs yielded a prominent env/rev amplification from 8 HIV-1 infected samples (**Fig. 2e**), randomly selected from a recent cohort of HIV-1 infected infants (45). Despite employing optimized primers selected from an exhaustive panel, the cloning efficiency of the env/rev cassettes via SGA amplification varied between samples. Further optimization of PCR for amplifying env/rev cassettes was done by varying the concentration of existing reactants s (magnesium, DMSO, BSA, PEG8000, primers). In addition, reducing the number of cycles in the first round of nested PCR significantly reduced spurious, non-specific products observed in certain samples.

**Table 1.**
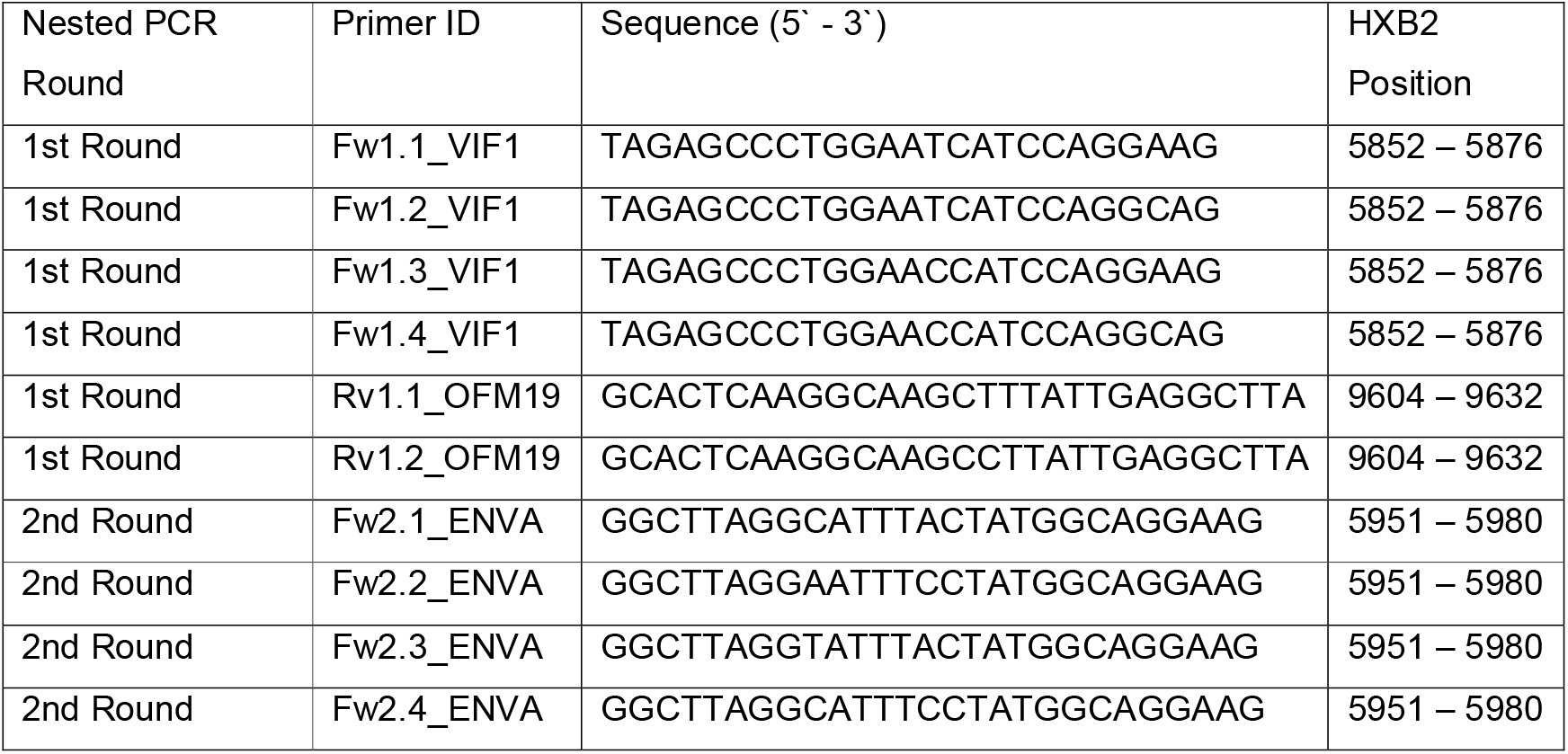
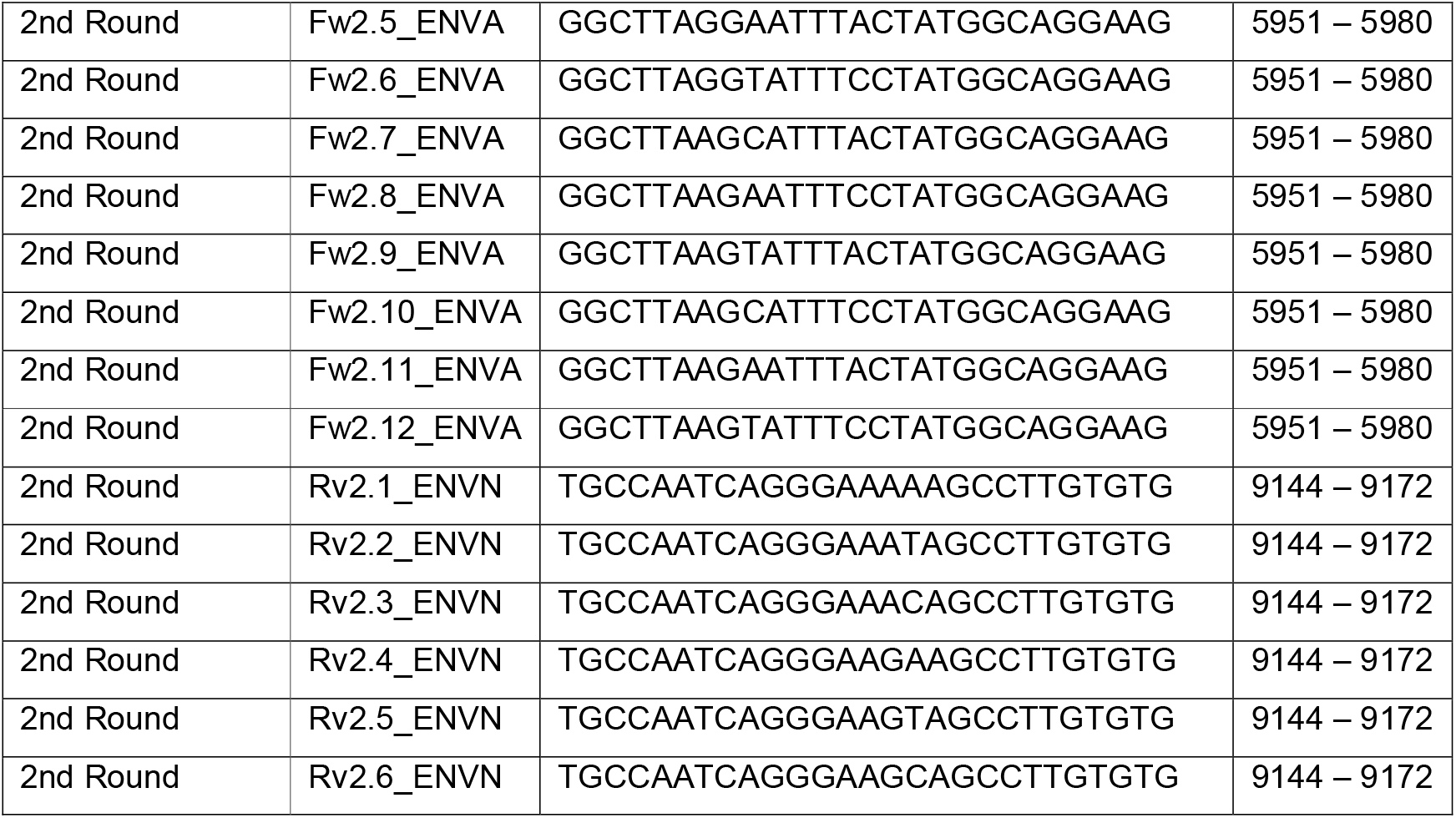
Primer pairs used for the amplification of HIV-1 Indian clade C env/rev cassettes. Primer pair Fw1.3_VIF1/Rv1.1_OFM19 for 1^st^ round PCR and Fw2.6_ENVA/Rv2.4_ENVN for 2^nd^ round gave most satisfactory env/rev cassette amplification across samples.

**Figure 1.**
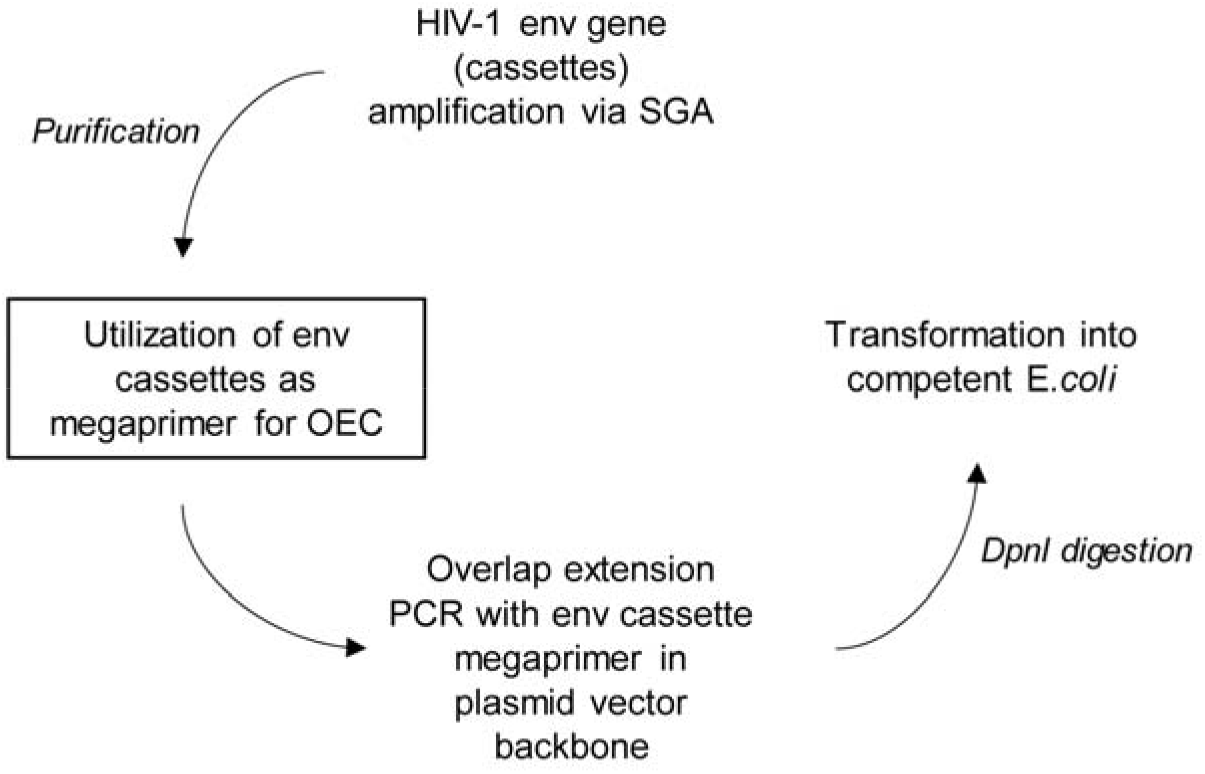
Schematic flow for overlap extension cloning. Outline of the major steps involved in OEC is provided. The key step outlining the utilization of env/rev cassettes as megaprimer is boxed. First, env/rev cassettes are PCR amplified from HIV-1 infected individuals and then used as megaprimers in plasmid vectors containing the integration sites (sequence of forward and reverse primer for env/rev cassette amplification) (see the vector map provided in supplementary figure 2), ultimately leading to generation of double nicked plasmid which is transformed into *E.coli*.

**Figure 2.**
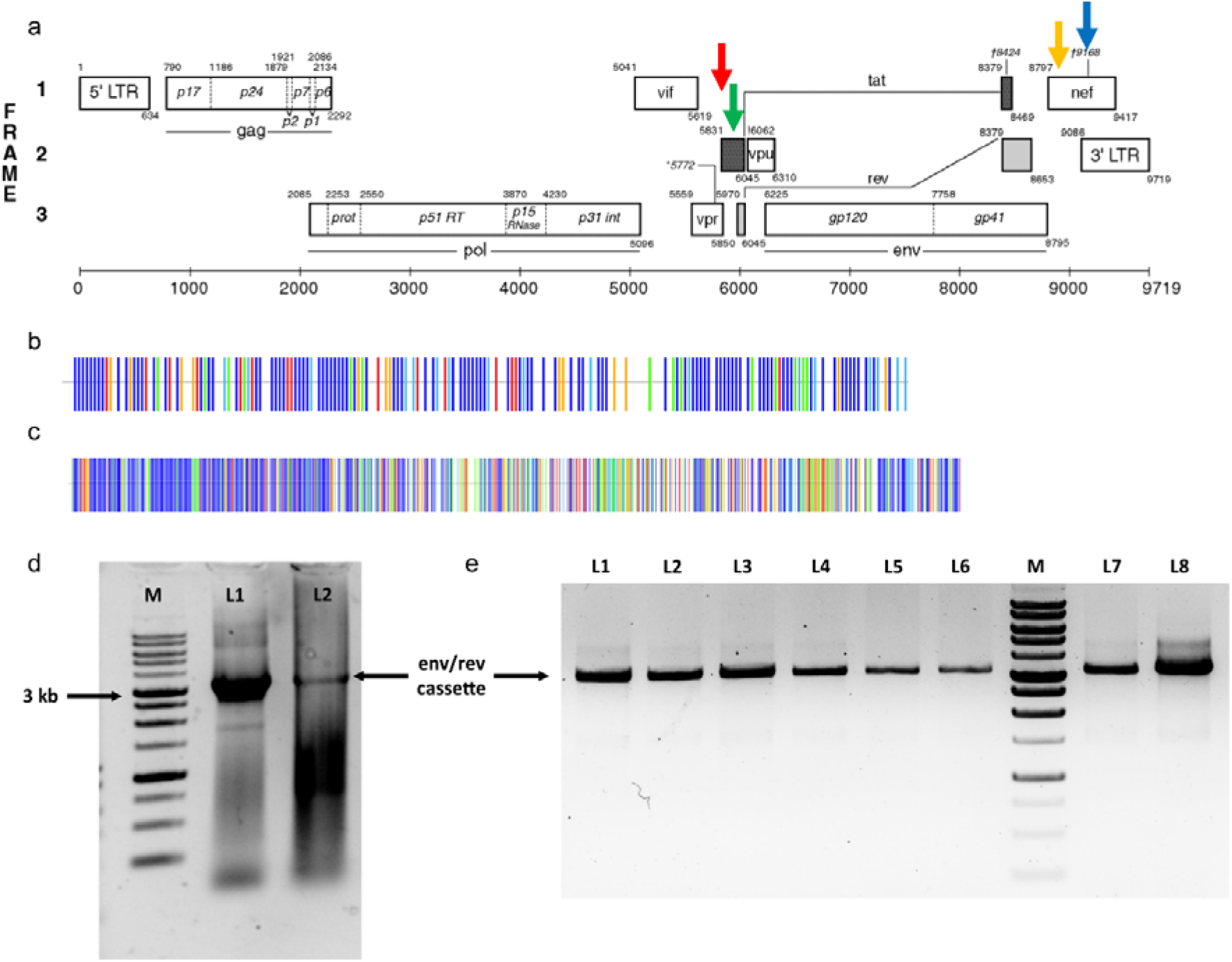
Optimizing the amplification of env/rev cassettes. (a) Schematic map of the HIV-1 genome showing the position of nested PCR primers. Within the first exon of tat, 1st round forward primer (Fw1) is marked by red, 2nd round forward primer (Fw2) is marked by green, and within the nef region 1st round reverse primer (Rv1) is marked by blue and 2nd round reverse primer (Rv2) is marked by yellow arrow. (b – c) Schematic map of the tat and nef region representing the sequence diversity in Indian clade C sequences reported in LANL HIV-1 database. Dark blue lines represent position that contain any of the four bases, while green lines represent adenine, red line represent thymine, orange line represent guanine and sky-blue lines represent cytosine relative to the Indian clade C consensus sequence generated by aligning available sequence from HIV-1 database. (d – e) Gel pictures showing amplified env/rev cassettes (~3.2 kb) from updated primer pairs as mentioned in table 1. In d, L1 represent plasmid control (HIV-25710, NIH AIDS Reagent Program #11505) while L2 represents amplified env/rev cassette using multiplex approach while in e, L1 to L8 represent env/rev amplification after optimization of primer binding using iterative selection approach.

To generate overlaps (plasmid sequences at both ends) that can be used for OEC, we added overhangs of 15, 25 and 35 bp randomly selected, upstream of the multiple cloning site (MCS) region of pcDNA3.1, to the chimeric primers generated by combining four primers, with 5` end complementary to the vector and 3` ends optimum for amplification of env/rev cassettes. Interestingly, none of the chimeric primer pair combination could amplify env/rev cassettes from the eight patient samples but prominent amplification was seen when plasmids containing env/rev cassettes were used. Weak amplification with 15 bp overhang chimeric primers were however observed in two of the eight samples (data not shown). HIV-1 contains an above average percentage of adenine (A) nucleotides, while cysteine (C) nucleotides are extremely low (**Supplementary Fig. 1**). Designing primers with standard primer designing rules to accommodate high HIV-1 diversity is therefore difficult. Primers used for amplification of env/rev cassettes are typically AT-rich while the overhang sequences taken from plasmids show typical pattern of base distribution (optimally dispersed pyrimidine and purine patterns). Addition of plasmid sequences to primers optimized for HIV-1 env/rev amplification, therefore, generates suboptimal chimeric primers (uneven distribution of pyrimidine and purine bases) plausibly explaining the failure of chimeric primers to amplify env/rev cassettes from biological samples. As amplification of env/rev cassettes with chimeric primers gave unsatisfactory results, we utilized an alternate approach for performing OEC with env/rev cassettes serving as megaprimers. We placed the Fw2 and Rv2 sequences 150 bp apart around the multiple cloning site (MCS) of pcDNA3.1 (pcDNA3.1_ITR), generating a plasmid that can be directly used for OEC using env/rev cassettes amplified from patient samples (**Supplementary Fig. 2**).

OEC has been reported to use linear amplification, typically generating amplicons too little to be visualized on agarose gels. In addition, the effectiveness of OEC has been shown to be dependent upon megaprimer concentration, with the megaprimer concentration being inversely proportional to size (26). To obtain good yield, four parameters were sequentially optimized, namely the number of cycles, annealing temperatures, concentration of recipient plasmid (pcDNA3.1_ITR), and concentration of megaprimer (env/rev cassette). We began our iterative optimization of OEC using 10 to 100 ng of pcDNA3.1_ITR and PCR purified env/rev cassettes at excess ratios of 1:3 to 1:300 in 25-μl reactions using previously reported reaction conditions. The PCR reaction was inhibited at high DNA concentrations (100 ng of pcDNA3.1_ITR and 1745 ng of PCR purified env/rev cassettes at 1:35 molar excess ratios) (**Fig. 3a**). High concentrations of insert and lower annealing temperature for megaprimers have been implicated as necessary for successful OEC (26,27), however we found that a high concentration of the env gene insert often led to failed PCR. A molar excess ratio of 1:5 to 1:10 of env/rev cassettes to 50 ng of pcDNA3.1_ITR gave the most optimum results, though multiple bands after OEC were observed, when the vector to insert ratio exceeded 1:25 (**data not shown**). Higher annealing temperatures were required to maximize OEC efficiency (**Fig. 3b**), plausibly by minimizing the formation of interfering secondary structures, typical for lentiviruses (42,43), as hairpin structures within the template region can hinder the PCR amplification. Addition of BSA and removal of DMSO serendipitously eliminated the non-specific multiple bands observed with OEC. As DMSO disrupts secondary structure formation, the observed decrease in spurious PCR amplification in its absence is counterintuitive, and requires further exploration. In addition, the high concentration of pcDNA3.1_ITR (10 ng) made DpnI digestion necessary in the OEC workflow, though DpnI could be added straight to Phusion HF buffer, thereby bypassing the need for purification steps. These reaction conditions were then taken forward for further optimization.

**Figure 3.**
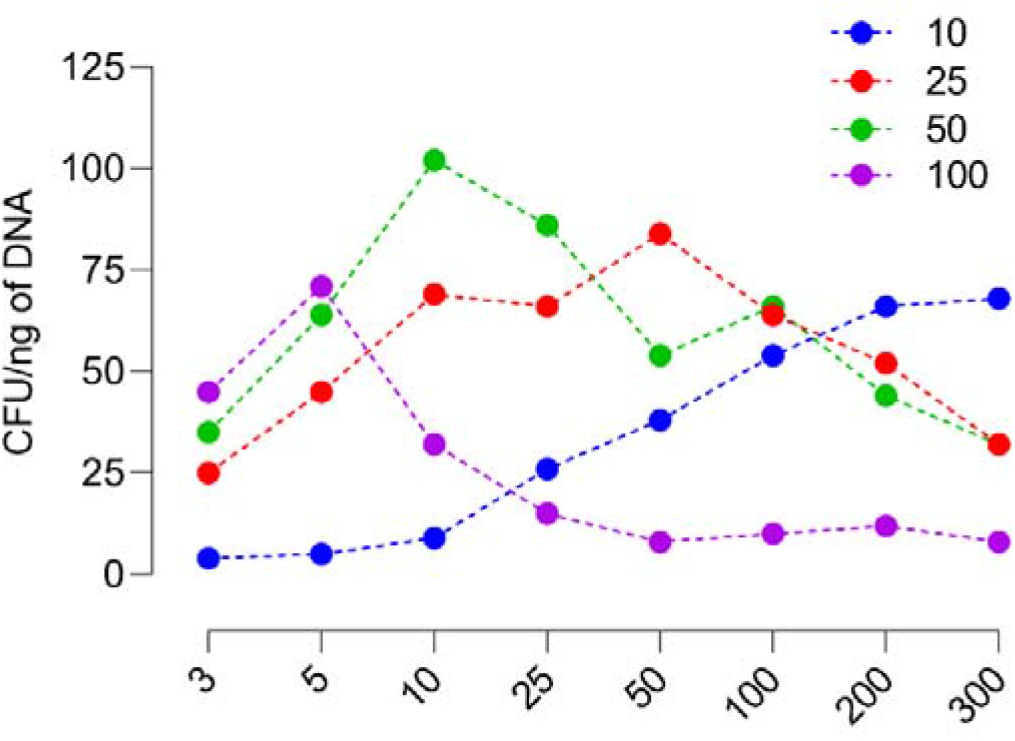
High concentration of megaprimer inhibits OEC. Overlap extension cloning of env/rev cassette from a single infant was cloned in pcDNA3.1_ITR at increasing ratios of vector to insert (1:3 to 1:300) as well as increasing amount of starting vector (10 to 100 ng of pcDNA3.1_ITR). Colony forming unit (CFU) were counted as the number of colonies normalized to starting vector amount. A vector to insert ratio of 1: 10 to 1:20 with starting vector amount of 10 ng showed maximum cloning efficiency across samples.

The number of cycles in OEC have been inversely linked to cloning efficiency and therefore to optimize the cycle number for amplification, we next performed OEC in increment of 2 cycles, with a minimum amplification of 10 cycles to begin with. Considerable variation was observed for OEC efficiency as a measure of PCR cycles between samples (cfu/ng of DNA, figure), though 14 – 16 cycles gave most satisfactory results (**Fig. 4a**). Few empty colonies were also observed in OEC done with pcDNA3.1_ITR alone (serving as the vector control), presumably due to the carry-over of the vector, as DpnI digestion was performed directly in PCR buffer. Though commercially available restriction enzymes are active in PCR buffers, reaction efficiency varies between samples. To address the unwanted vector carryover, we subcloned CCDB gene (46) (from vector pZM247Fv1∆Env, procured from NIH AIDS Reagent Program) in-frame to T7 Promoter and between the Fw2 and Rv2 sites of pcDNA3.1_ITR via OEC. Utilization of the suicide vector (pcDNA3.1_ITR_CCDB) containing CCDB gene for OEC of env/rev cassettes completely removed unwanted vector carry-over as well as the requirement for DpnI digestion.

**Figure 4.**
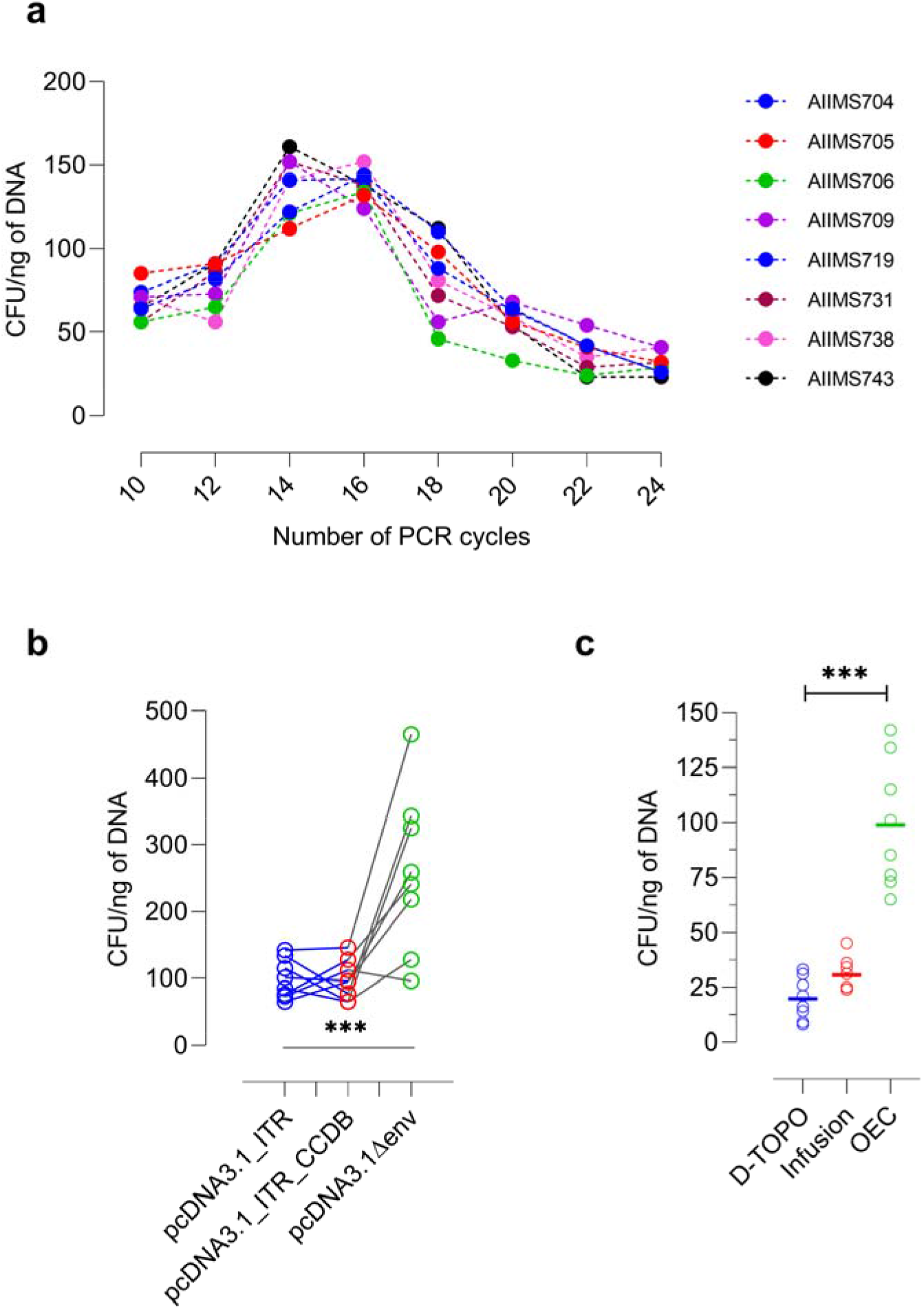
OEC efficiency as a function of amplification cycle. Overlap extension cloning performed for env/rev cassettes amplified from eight infant plasma samples was iteratively optimized as a function of amplification cycles in OEC PCR. 14 – 16 cycles consistent gave satisfactory cloning efficiency across samples. Colony forming unit (CFU) were counted as the number of colonies normalized to starting vector amount. (b) Diverse vector backbones were utilized to further improve the OEC cloning efficiency. Using pcDNA3.1∆env vector backbone, maximum efficiency of OEC was achieved. (c) Comparison of the cloning efficiency of the topoisomerase mediated (D-TOPO), T4 DNA polymerase mediated (Infusion) and overlap extension PCR mediated (OEC) cloning strategies for cloning the env/rev cassettes from eight HIV-1 infected infants.

Considering our initial goal was to develop a system that can be globally implemented across labs working with HIV-1, we reasoned using readily available pseudoviral env clones used across labs for standardized assessment of bnAbs or vaccine-induced sera nAbs as vectors for OEC. Herein, we first mutated the ENVA and ENVN sites in 329_14_B1 backbone (a subtype C pseudoviral clone, accession number MK076593) to our primer pairs (**refer to Table 1**), added a stop codon prior to V1 region and used it as a vector for cloning env/rev cassettes amplified from patient samples via OEC. Addition of stop codon was necessary as it simplified the identification of functional clones for further downstream processing, given colony PCR was not possible as OEC was used to swap a 3.2 kb fragment with a similar sized env/rev cassette from biological samples. Interestingly, using this pcDNA3.1 delta env backbone gave significantly higher number of colonies compared to colonies observed with either pcDNA3.1_ITR or pcDNA3.1_ITR_CCDB suicide vector (**Fig. 4b**), presumably due to increased regions of homology upstream to the ENVN and downstream to the ENVA ITR sites in 329_14_B1∆Env. To compare the efficiency of OEC, we then cloned the respective env/rev cassettes from the eight infant samples using the commercially available topoisomerase mediated (TOPO Cloning) and T4 DNA polymerase mediated (Infusion Cloning) cloning kits, and observed significantly higher cloning efficiency, indicating that OEC can be used to efficiently clone difficult-to-clone HIV-1 env gene (**Fig. 4c**)

Furthermore, we attempted to simplify and standardize the OEC protocol into a one-pot (single tube) approach for cloning env/rev cassettes. Using the unpurified PCR amplified env/rev cassettes from a sample that gave negligible non-specific background, we could successfully perform OEC as a one-pot platform for cloning env/rev cassettes, though a significantly high number of colonies containing the unwanted primer-dimers as insert were observed. Further titrating the initial concentration of primers used for env/rev amplification did decrease the number of unwanted primer-dimer colonies, but unwanted clones containing primer-dimers could not be completely eliminated. Utilizing touchdown PCR for amplification of env/rev cassettes minimized the number of spurious primer-dimer clones to almost negligible, though it also led to reduced number of colonies after successful OEC. Of particular note, one-pot approach only worked for env/rev cassettes amplified via SGA and not bulk PCR. Amplification of env/rev cassettes with touchdown PCR, followed by one-pot OEC reaction using higher amount of vector, led to the most satisfactory results, though for better performance, PCR purification prior to OEC is necessary. Touchdown PCR (TD-PCR) was developed as a simpler solution to address the mispriming and production of non-specific (47), spurious products that often result due to poor melting temperature estimations and improper annealing conditions but in case of env/rev cassette amplification from biological samples, even the most optimized PCR conditions gave non-specific amplification in certain samples. PCRs for env/rev cassettes require optimization for each biological sample but given the extensive diversity and circulating mutant pool in an infected individual, optimized primers, on average, can minimize the requirement for PCR optimization from sample to sample.

Once we successfully swapped patient amplified env/rev cassettes into the HIV-25710_2_42 backbone, another versatility of OEC became apparent. Given that swapping env/rev cassettes from one source into another could easily be achieved via OEC, we next utilized the same procedure to generate chimeras between different env/rev cassettes. Phusion polymerase has been reported to fail PCR amplification when complementary Fw and Rv primers (as suggested for QuikChange Kits) are used for site-directed mutagenesis (48). In our case, we observed persistent positive amplification when using Phusion to perform OEC for site-directed mutagenesis or chimeragenesis, though the primers had to be on average of 60-65 bp long to achieve more than 90% efficiency via OEC. For chimeragenesis, primers longer than 25 bp were required for efficiencies >90%. Though, of note, if DpnI digestion is optimum, OEC is an all or none approach, thereby, with reaction efficiencies approaching 100%. As noted for several other cloning approaches, and reported previously for OEC, cloning efficiency of OEC in the context of HIV-1 mutagenesis and chimeragenesis was a function of insert length, with considerably lower number of colonies observed for larger inserts (swaps for chimeras) compared to smaller inserts (**Fig. 5**).

**Figure 5.**
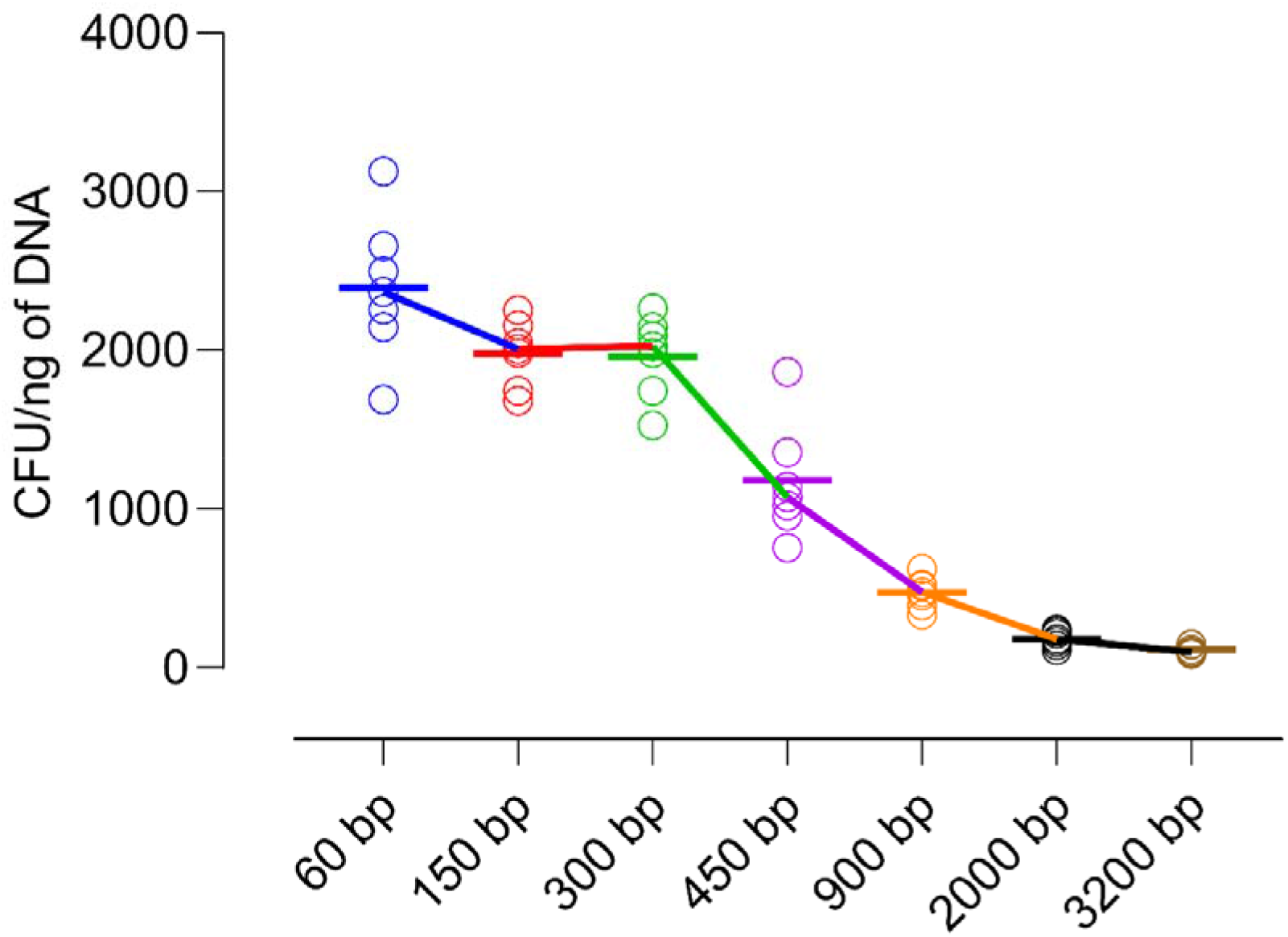
Efficiency of Overlap Extension Cloning for chimeragenesis in env/rev cassettes. With varying length of megaprimers (inserts to be swapped), Overlap extension cloning was performed for chimeragenesis. Results are reported for six independent experiments. Colony forming unit (CFU) were counted as the number of colonies normalized to starting vector amount. All the megaprimers were swapped into the vector backbone pcDNA3.1-329_14_B1. Marked reduction in cloning efficiency (chimeragenesis) with increasing length of insert was observed though satisfactory results were still observed.

Cloning of HIV-1 env/rev cassettes via OEC is easy to monitor and can be optimized for distinct biological samples with minimum modifications to the final protocol described in this study. For OEC, general rules for primer design apply, but additional focus is required for both the 5` - and 3` - ends with even distribution of pyrimidine and purine bases and either a G or C at the terminus. Under suboptimal conditions, OEC significantly generates unwanted clones, and therefore, optimization of reaction conditions is a necessity. Utilizing touchdown PCR further increases the efficiency of OEC for cloning env/rev cassettes as higher annealing temperatures compared with the expected primer Ta during the initial PCR cycles results in increased PCR product specificity, as spurious primer-template interactions are less stable than the specific ones. Substantial non-specific amplification often necessitates PCR purification prior to OEC, and utilization of TD-PCR further simplifies the OEC workflow. To validate the optimized OEC strategy, HIV-1 env/rev cassettes were amplified from 8 infected infants, of which only, a single sample required modified protocols (varied Ta) for successful OEC. The flexibility and high-throughput of OEC allows far greater versatility than the current global standard of TOPO mediated cloning for characterizing HIV-1 envelope glycoprotein. SGA amplified env/rev cassettes could be integrated into the vector by a single PCR, bypassing the need for utilizing expensive cloning kits, or novel enzymes such as T4 DNA polymerase, Taq DNA ligase, exonucleases and recombinases. The requirement to continuously monitor and optimize OEC is in parallel to any long PCR protocol, and requires moderate level of experience and understanding on the user’s part. To summarize, herein, we report OEC as an alternate PCR-based approach for cloning HIV-1 env/rev cassettes, that can yield high-efficiency amplification of HIV-1 envelopes with diverse sequence variability.

## Materials and Methods

### Materials

Phusion DNA polymerase, dNTPs, DpnI, DMSO and BSA were purchased from Thermo Scientific. *E. coli* TOP10 (Invitrogen) was used for cloning and competent cells were prepared in-house (10^6^ – 10^7^ cfu/μg). DNA oligonucleotides were synthesized by Eurofins genomics, India. The pcDNA3.1 (+) vector was procured from Invitrogen. The pZM247Fv1∆Env was acquired from NIH AIDS Reagent Program (#11940). The lab generated pseudovirus 329_14_B1 is described previously(18).

### Designing modified primers for env/rev amplification using HIV-1 Alignments

The Indian Clade C tat and nef sequences were compiled from the Los Alamos National Laboratory HIV-1 Sequence Database using the sequence search interface (https://www.hiv.lanl.gov/components/sequence/HIV/search/search.html), and codon-aligned using the HIV align (https://www.hiv.lanl.gov/content/sequence/VIRALIGN/viralign.html) tool using the HMM-align model with compensating mutation occurring within 5 codons to compensate for frameshift. The MEGA-X and ClustalX2 software tools were used for manually editing the alignments. Schematic plots to represent the sequence diversity, a measure of mismatches in the alignments, were generated using the highlighter tool (https://www.hiv.lanl.gov/content/sequence/HIGHLIGHT/highlighter_top.html) with sequences sorted by similarity and gaps in the alignment treated as characters. Previously described primers were used as anchors to design optimized primers for Indian Clade C sequences. Overlaps comprising of 15, 25, and 35 base pairs [5’ to the BamHI site, and 3’ to the ApaI site in pcDNA3.1(+)] with the primer pairs were manually incorporated using the SnapGene Viewer Program. Primers for site-directed mutagenesis of the pseudovirus 329_14_B1 were designed using Agilent’s quikchange primer designing tool. Primers for chimeragenesis were manually designed based on swapping inserts of varying length into the 329_14_B1 HIV-1 pseudoviral clone utilizing SnapGene Viewer Program, with normal primer designing rules of 20-25 bp length with a Tm of approximately 60°C, ideally ending with G/C with last 6 bp containing equal number of pyrimidine and purine bases.

### Vector design and construction

pcDNA3.1_ITR, pcDNA3.1_ITR_CCDB and pcDNA3.1delta Env were engineered using the mammalian expression vector pcDNA3.1 (+) through a series of overlap extension cloning reactions. pcDNA3.1_ITR was generated via cloning ITR sites ENVA (5` - CACCGGCTTAGGAATTTACTATGGCAGGAAG - 3`) and ENVN (5` - TGCCAATCAGGGAAAAAGCCTTGTGTG - 3`) upstream to BamHI and downstream to ApaI restriction site respectively. Vector pcDNA3.1_ITR_CCDB, CCDB gene was PCR amplified from vector ZM247Fv1∆Env using primers ENVA_CCDB_Forward (5` - TAGGAATTTACTATGGCAGGAAGATGCAGTTTAAGGTTTACACC - 3`) and ENVN_CCDB_Reverse (5` - GCCAATCAGGGAAAAAGCCTTGTGTGTTATATTCCCCAGAACATCAGGTTAATGG - 3`). The amplicon was next swapped using OEC into 10 ng of vector pcDNA3.1_ITR using CCDB gene as megaprimer at a ratio of 1:350. The PCR conditions used were an initial denaturation at 98°C for 2 min followed by 16 cycles of denaturation at 98°C for 10 sec, annealing at 64°C for 30 sec, and extension at 72°C for 3 min and a final extension at 72°C for 5 min. The pcDNA3.1∆Env (in pseudoviral backbone 329_14_B1) was generated by inserting a stop codon prior to V1 region of the env gene using primers V1_Stop_Forward (5` - CAGATGCAGGAGGATGTAATCAGTTTAATGGGATCAAAGCCTAAAGCCATGTG - 3`) and V1_Stop_Reverse (5` - CACATGGCTTTAGGCTTTGATCCCATTAAACTGATTACATCCTCCTGCATCTG - 3`). All transformation reactions were performed using in-house prepared TOP10 competent cells followed by sequencing to confirm successful generation of vectors.

### Amplification of env/rev cassettes and overlap extension cloning (OEC) PCR

The env/rev cassettes of HIV-1 from infected infants were amplified as described previously. Briefly, viral RNA was isolated from 140 μl of plasma using QIAamp Viral RNA Mini Kit, reverse transcribed, using gene specific primer OFM19 (5` - GCACTCAAGGCAAGCTTTATTGAGGCTTA – 3`) and Superscript IV reverse transcriptase into cDNA. The cDNA was then used as template to amplify the envelope gene using High Fidelity Phusion DNA Polymerase (New England Biolabs). Nested PCRs were performed, using primers given in table 1 with the following PCR conditions; an initial denaturation at 98°C for 2 min, followed by 25 cycles of denaturation at 98°C for 10 sec, annealing at 59°C for 30 sec, and extension at 72°C for 2 min and a final extension at 72°C for 5 min. A second round PCR was performed using first round amplicons with reaction conditions of initial denaturation of 98°C for 2 min, followed by 35 cycles of denaturation at 98°C, annealing at 59°C for 30 sec, and extension at 72°C for 90 sec, and a final extension at 72°C for 90 secs. The env/rev cassettes were PCR purified using the QIAquick PCR & Gel Cleanup Kit and utilized as megaprimers for overlap extension cloning into pcDNA3.1 vector backbone (pcDNA3.1_ITR, pcDNA3.1_ITR_CCDB, and pcDNA3.1∆Env). Overlap extension cloning was performed with varying amount of pcDNA vector backbone and env/rev cassettes with PCR reaction conditions of initial denaturation at 98°C for 2 min, followed by 10 – 24 cycles of denaturation at 98°C for 10 sec, annealing at 62°C for 30 sec, and extension at 72°C for 4.5 min, and a final extension of 5 min at 72°C. In addition, 1% DMSO and 1% BSA (1 mg/ml) were used as additives for OEC of env/rev cassettes. The PCR products were analyzed using 0.8% Agarose gel electrophoresis.

### Transformation and DNA Sequencing

Aliquots (5 μl) of undigested and DpnI digested OEC PCR amplicons were directly transformed into in-house generated chemically competent TOP10 cells. Cells were spread on 2XYT-agar plates containing 100 μg/ml of ampicillin sodium salt and incubated overnight at 30°C. For env/rev cassettes, incubation at lower temperature reduced the toxicity associated with problematic DNA sequences (42–44). The number of colonies grown on each plate were calculated and normalized to per ng of starting vector backbone to calculate cloning efficiency (colony forming units per ng of DNA). DNA Sequencing was performed commercially from eurofins genomics, India.

## Acknowledgments

We are thankful to all the study subjects for participating in this study and the NIH AIDS Reagent program for providing HIV-1 envelope pseudovirus plasmids.

## Author Contributions

N.M, S.S and K.L designed the study. N.M, A.D and S.S performed the experimental work. N.M analyzed the data, wrote the initial manuscript, revised and finalized the manuscript. K.L wrote, edited, revised and finalized the manuscript.

## Data and Material Availability

All data required to state the conclusions in the paper are present in the paper and/or the supplementary data. Additional information related to the paper, if required, can be requested from the authors.

## Supplementary Figures

**Supplementary Figure 1.**
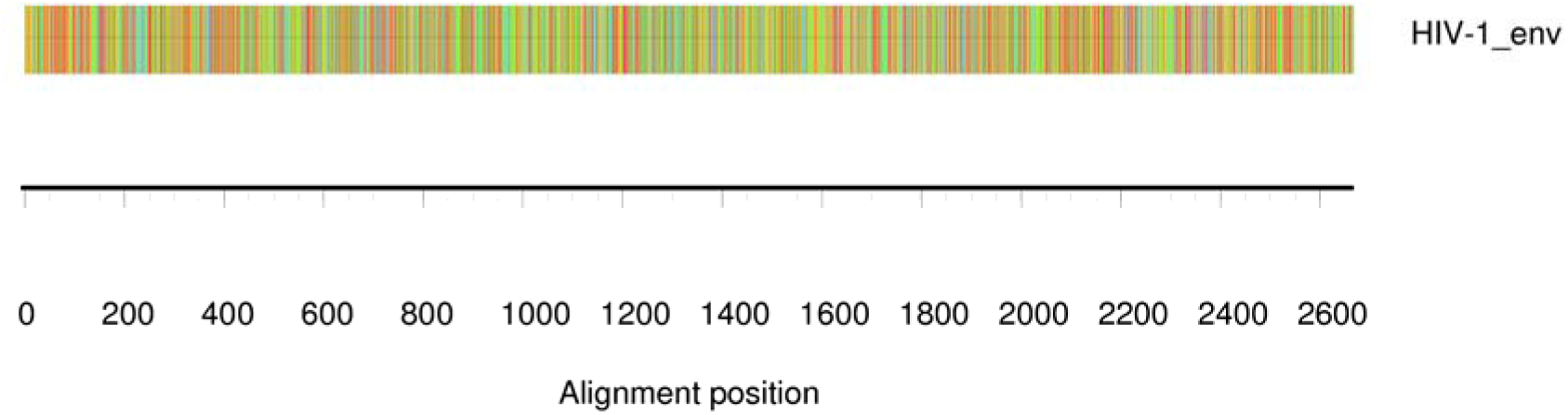
Unusual base composition of HIV-1 env gene. HIV-1 contains an above average percentage of adenine (A) nucleotides, while cysteine (C) nucleotides are extremely low as evident by the large percentage of red (adenine), and limited percentage of blue (cytosine) lines in the base composition plot for consensus env sequence (HIV-1_env) generated by aligning the env sequences of Indian clade C available in LANL HIV-1 database https://www.hiv.lanl.gov/components/sequence/HIV/search/search.html).

**Supplementary Figure 2.**
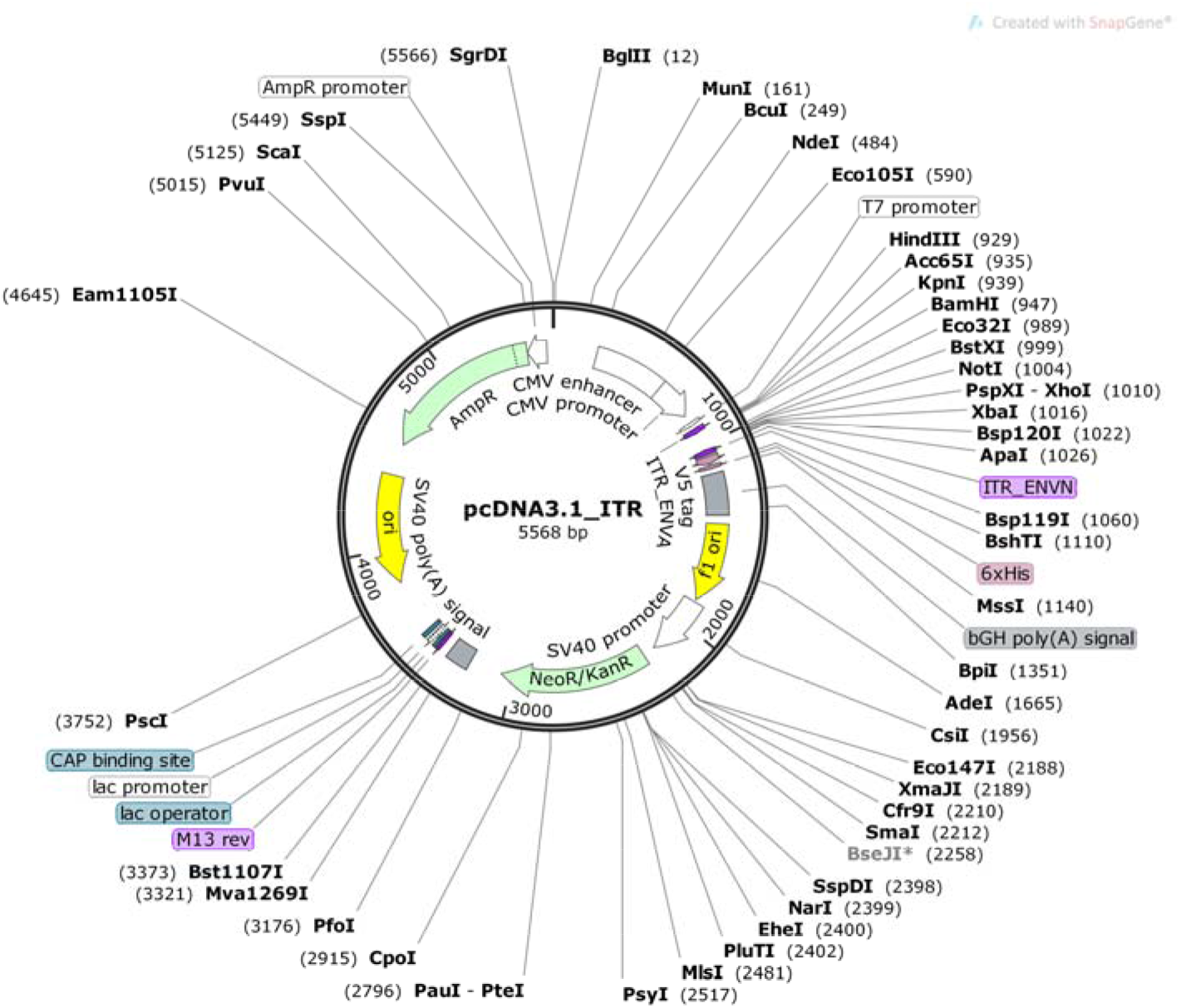
Vector map of modified pcDNA3.1 plasmid utilized for cloning HIV-1 env/rev cassettes. In pcDNA3.1 (+) backbone, ENVA (Fw2) and ENVN (Rv2) sites optimized based on results of figure 1 were integrated (ITR_ENVA and ITR_ENVN) into pcDNA3.1 by performing two rounds of overlap extension PCR upstream to HindIII and downstream to ApaI restriction sites.

